# Subtype-specific neutralizing antibodies promote antigenic shift during influenza virus co-infection

**DOI:** 10.1101/2025.08.08.669311

**Authors:** Mengnan Liu, Xiaoran Gao, Xueyang Li, Jinyang Hao, Christine Light, Yu Tian, Siqin Wang, Wenyu Yan, Rong Hai, Wei Hu, Guojun Wang

## Abstract

Reassortment between different influenza strains occurs when they co-infect the same host cell. The emergence of a reassortant virus depends on both its intrinsic fitness and extrinsic factors, including pre-existing humoral immunity. The generation of pandemic strains, such as H2N2 and H3N2, and zoonotic influenza A viruses, like H5N6, H5N8, and H7N9, in birds is suggested to be the result of extensive selection by pre-existing antibodies. To further explore the role of humoral immunity in reassortment, we generated two divergent fluorescent protein-expressing viruses and used strain-specific and cross-reactive monoclonal antibodies (mAbs) to assess the impact of cross-immunity on reassortment. Our results indicate that all mAbs altered the genotypic diversity and significantly reduced the release of progeny virions in co-infected cells both *in vitro* and *in vivo*. Moreover, antibody transfer studies in mice revealed protection from challenge with divergent pathogenicity profiles. Notably, selection driven by a strain-specific mAb depended on its neutralizing specificity, whereas the selection driven by broadly reactive mAbs was independent of neutralization specificity. Our findings demonstrate that pre-existing neutralizing antibodies shape reassortment and that strain-specific neutralizing antibodies promote antigenic shift during co-infection, which is not the case for broadly cross-reactive antibodies that recognize influenza viruses from different subtypes.

## 1. Introduction

Influenza viruses are highly variable pathogens that uses two mechanisms of antigenic variation to evade the host immune system: antigenic drift and antigenic shift in its hemagglutinin (HA) and neuraminidase (NA) genes [1–3]. Antigenic drift, which arises from the error-prone nature of RNA-dependent RNA polymerase, allows the virus to generate escape mutants that can evade recognition by pre-existing antibodies, thereby facilitating localized influenza outbreaks [4,5]. In contrast, antigenic shift involves a more drastic change in HA and/or NA genes, which can occur through reassortment between human and nonhuman viruses or through host switching from animals to humans [1–3]. The introduction of novel antigens resulting from antigenic shift enables the virus to spread rapidly in a naive population and has historically been associated with pandemics, as exemplified by the outbreaks of 1918 (H1N1) [4,5], 1957 (H2N2) [6], 1968 (H3N2) [7], and 2009 (pH1N1) [8], all of which involved reassortment.

The segmented nature of the influenza virus genome allows for reassortment to occur when two or more genetically distinct viruses co-infect the same host cell [9,10]. This process, theoretically, can generate 256 possible genotypes from two parental influenza A viruses (IAVs) [11]. However, current evidence suggests that only a limited subset of progeny combinations are produced, implying that the formation and propagation of reassortant progenies are biased [12,13]. The frequency of reassortment is influenced by several factors, including the co-infection of the same cell by both viruses [9,14]. This requires that both viruses have the ability to infect the same cell, a process governed by factors such as viral prevalence, timing and dose of infection, spatial dynamics of viruses in the host, and superinfection interference [9],[15–18]. Furthermore, accumulating evidence indicates that the association of influenza genes during reassortment is selective rather than random, with specific incompatibilities between heterologous viral components. These incompatibilities, referred to as segment mismatches, can be categorized as RNA-based or protein-based. For example, sequence differences in packaging signals can lead to incompatibilities between heterologous viral RNAs (vRNAs) [19,20]. Similarly, mismatch between heterologous proteins, including polymerase subunits, HA/NA proteins, and NS-vRNP complexes, can significantly influence the outcomes of reassortment between divergent IAV strains [11,21–25].

In addition to the intrinsic factors that influence reassortment, extrinsic factors also play a significant role in suppressing this process. The influenza A virus (IAV) genome encodes 11 or more proteins, but two major surface glycoproteins, hemagglutinin (HA) and neuraminidase (NA), are of particular importance due to their immunogenicity [26,27]. HA is characterized by a highly variable head domain and a relatively conserved stalk domain [28]. Antibodies that bind to the head domain of HA are immunodominant but often strain-specific [27], whereas antibodies that target the stalk domain are less abundant and less potent, yet cross-reactive across a broad range of virus isolates and subtypes [29–32]. Notably, antibodies towards NA are immunosubdominant and exhibit direct antiviral activity [33,34]. Both anti-HA and anti-NA antibodies provide protection in animal models through passive transfer experiments [29,33]. As HA and NA play critical roles in different stages of the virus life cycle, antibodies against these proteins exhibit distinct antiviral mechanisms. Despite significant advances in understanding the humoral immune response to IAV, the impact of antibodies on reassortment remains poorly understood [28,29,33,35].

In this study, we first generated two antigenically diverse reporter viruses: a mCherry-expressing A/PuertoRico/8/34 (PR8-mCherry) H1N1 virus and a green fluorescent protein (GFP)-expressing A/Viet Nam/1203/2004 (VN-GFP) H5N1virus. We then developed a single-cell sorting assay to quantify reassortment and assess the impact of antibodies on this process. Three mAbs specific to the HA of the parental viruses were chosen to examine their effects on reassortment. These mAbs exhibited neutralizing activity against both parental viruses in co-infected cells, both *in vitro* and *in vivo*. Notably, the strain-specific neutralizing mAbs altered the reassortment outcomes by selectively inhibiting the production of progeny virions bearing the neutralized HA, while promoting the generation of virions with mismatched HA. These findings suggested that strain-specific neutralizing antibodies can modulate reassortment to drive antigenic shift towards antibody escape.

## 2. Results

### 2.1. Generation and characterization of fluorescent protein-expressing viruses

Using reporter viruses offers opportunities to identify and trace influenza virus infection. To facilitate the study of IAV reassortment, two replication-competent fluorescent viruses (PR8-mCherry and VN-GFP) were generated by inserting a fluorescent reporter gene into the NS gene segment (Fig. 1A) [36]. We successfully rescued the PR8-mCherry and VN-GFP viruses using the established reverse genetics system, which formed visible fluorescent-expressing plaques (Fig. 1B and C) [36,37]. To determine the virus fitness, we compared growth kinetics of the fluorescent and parental viruses in Madin-Darby canine kidney (MDCK) cells and mice. Even though PR8-mCherry showed a delay in replication kinetics in MDCK cells at both multiplicity of infection (MOI) of 1 and MOI of 0.001, it grew to titers reaching up to 5 × 10^7^ plaque-forming units (pfu)/mL and 1.45 × 10^9^ pfu/mL, respectively, while the growth kinetics of VN-GFP were indistinguishable from the parental strain (Fig. 1D). Similar to previous findings for the attenuation of PR8 expressing reporter genes [36,38], a pathogenicity analysis in mice revealed that the virulence of PR8-mCherry was substantially lower than that of PR8; the dose required to kill 50% of infected mice (MLD_50_) was 10^4.5^ pfu for PR8-mCherry compared with 10^2.5^ pfu for PR8. Similar to the *in vitro* result, the replication kinetics experiments *in vivo* revealed similar MLD_50_ values between VN-GFP and VN (10^5.5^ pfu & 10^5.38^ pfu) (Fig. 1E). To test the stability of fluorescent viruses, we passaged the viruses thrice in MDCK cells and mice and performed plaque assays using supernatants from infected MDCK cells and lung homogenate from infected mice. We detected high fluorescent expression levels during its replication *in vitro* and *in vivo* (Fig. 1F). This indicated that the genetic modifications to the IAV genome did not result in a loss in virus fitness, allowing for the assessment of co-infection and reassortment both *in vitro* and *in vivo*.

**Fig. 1.**
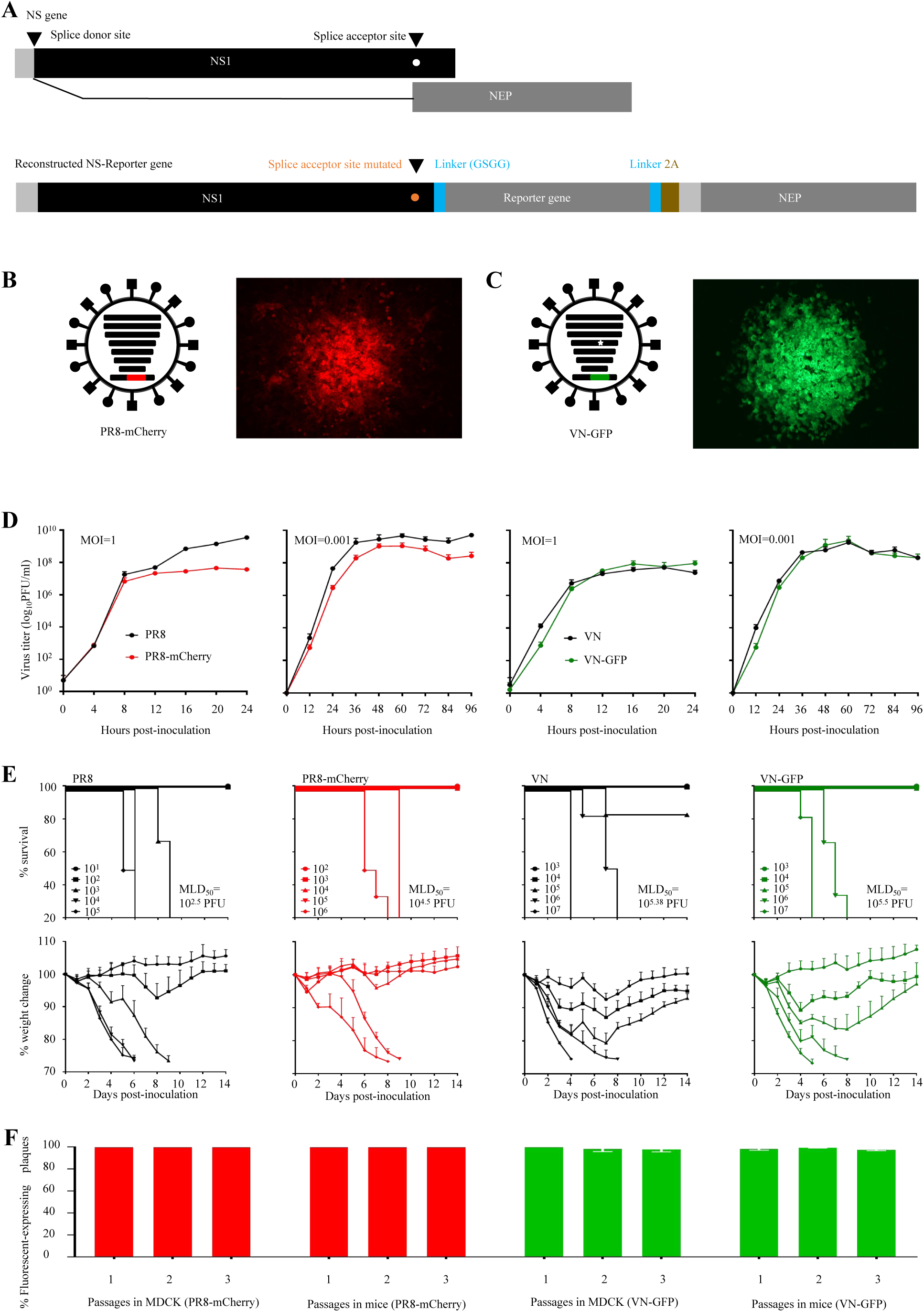
Generation and characterization of the fluorescent-expressing Viruses. (A) Schematic diagrams of the NS segment from wild-type (WT) virus and fluorescent-expressing virus. (Top) The WT NS segment expresses NS1/NEP proteins by mRNA splicing. (Bottom) The fluorescent-expressing NS expresses both NS1-reporter and NEP proteins from a single mRNA. Silent mutation was introduced into the splice acceptor site in NS1 to prevent mRNA splicing. NS1(black) was fused to the reporter gene (light gray) via a GSGG linker (blue), followed by the 2A autoproteolytic cleavage site (brown) and NEP (dark gray). (B and C) Fluorescent expression of the reporter viruses in MDCK cells. MDCK cells were infected with (B) PR8-mCherry or (C) VN-GFP, and at 48 h.p.i., the reporter expression of each virus plaque was observed by using fluorescence microscopy. The white star in panel C indicated removing of the multi-basic cleavage site of HA to reduce virulence. (D) Replication kinetics of WT and reporter viruses. MDCK cells were infected with the viruses at an MOI of 1 or 0.001, and culture supernatants were harvested at the indicated times and titrated by plaque assay. (E) Virulence of WT (PR8 and VN) and reporter (PR8-mCherry and VN-GFP) viruses in mice. Female 6-to-8-week-old BALB/c mice (n =5) were intranasally inoculated with indicated viruses and monitored daily for 2 weeks for body weight loss and survival. Mice that lost 25% or greater of their initial body weight were sacrificed. The MLD50 was determined by the method of Reed and Muench. (F) In vivo and in vitro stability of PR8-mCherry and VN-GFP. MDCK cells were infected with the reporter viruses at an MOI of 0.001. Supernantant was collected at 24 h.p.i.. Mice were infected with 10^4^ pfu of reporter viruses and the lung was collected at three d.p.i.. The percentage of fluorescent-expressing plaques in supernatants or lung homogenates was analyzed by plaque assay. The viruses were blindly passaged thrice in MDCK or mice. Each data point represents the mean ± deviation of replicates (n =3).

### 2.2. Single-cell sorting assay for genotyping fully infectious virions released from cells infected with parental viruses

Numerous studies have demonstrated that less than 10% of IAV particles released by infected cells are fully infectious [39,40]. Non-infectious virions can complement and initiate replication at high-multiplicity IAV infection, failing to cleanly separate non-infectious virions from infectious progenies in endpoint dilution assays (the 50% infectious dose [ID_50_] or plaque forming units [pfus]).

To ensure accurate genotyping of the infectious virions released from infected cells, we performed a single-cell sorting assay (Fig. 2A). First, a population of MDCK cells was infected with PR8-mCherry and VN-GFP at indicated MOIs. Twelve hours post-infection (h.p.i), the supernatants were collected for further analysis of the virions, and the cells were harvested to determine the infection rate of the parental viruses with flow cytometry. To avoid the co-infection of the progenies, the supernatants were inoculated into the MDCK cells with a low MOI (less than 1% of the cells were fluorescent protein-expressing). At six h.p.i., each fluorescent-expressing cell was sorted by Fluorescence-activated cell sorting (FACS) into a well containing mono-layer MDCK cells. The sorted cell infected with a fully infectious virion produced progeny, which could infect neighboring cells, and the well was entirely or partially fluorescent protein-expressing after 48 h of incubation. RNA from a fluorescent-expressing well was extracted and then reverse-transcribed into DNA with a universal primer. DNA melting peak profile-based high-resolution melting (MP-HRM) analysis was employed to genotype the fully infectious progenies. Briefly, the polymorphic region of each segment was used to implement the quantitative PCR (qPCR). The amplicon subjected to HRM analysis was used to distinguish the parental origin of each segment in each well based on the melting curve shape. The melting peak of each segment is shown in Figure 2B.

**Fig. 2.**
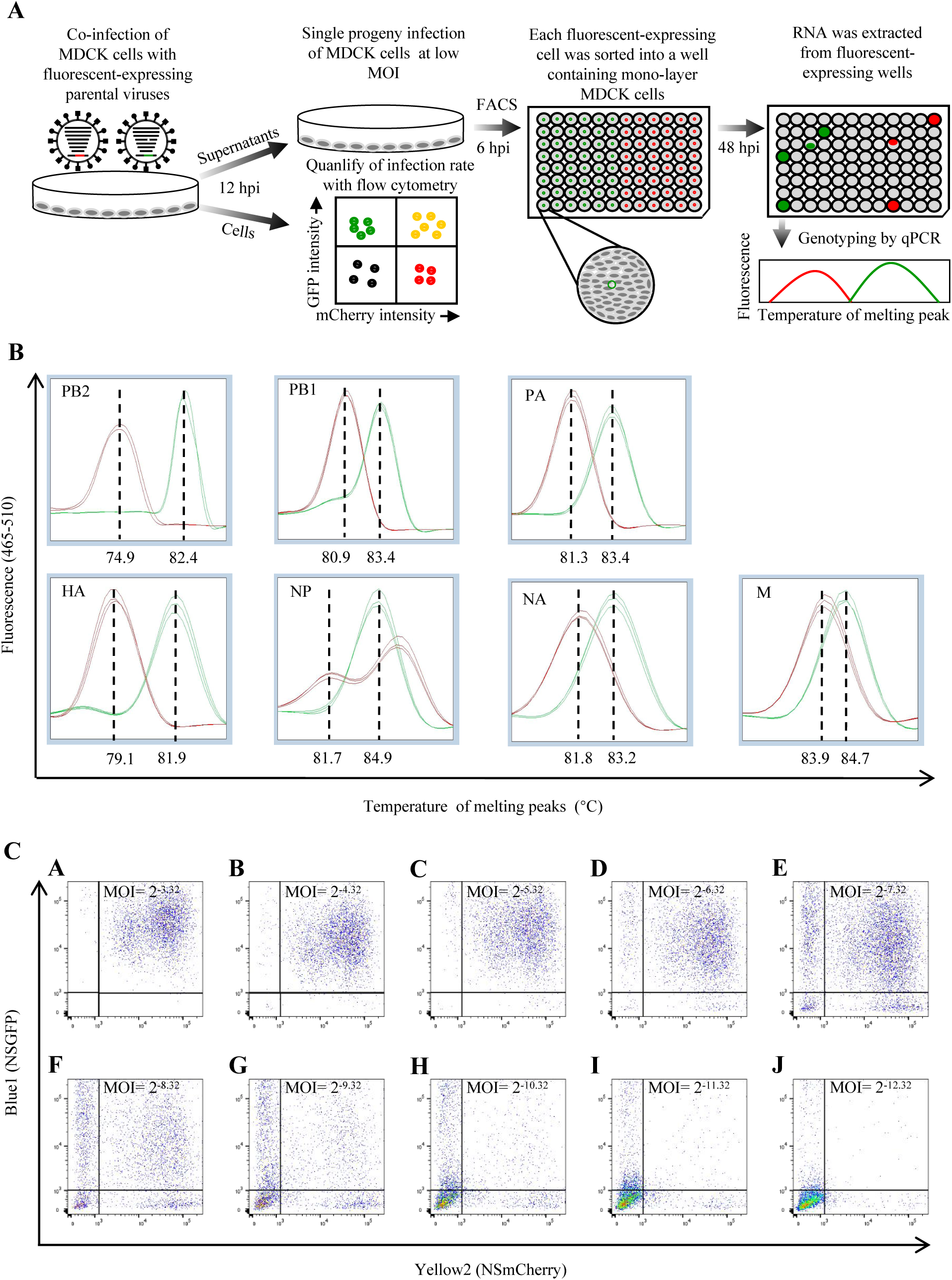
Experimental design to measure co-infection and determine virus genotype. (A) Schematic depicting our strategy for measuring the infection of parental viruses and genotyping the fully infectious virions with a single-cell sorting assay. MDCK cells were simultaneously infected with equal amounts of PR8-mCherry and VN-GFP viruses. At 12 h.p.i., supernatants were collected to genotype the released virions, and cells were harvested to determine the co-infection rate. MDCK cells were infected with supernatants at a low MOI. After six h of incubation, the cells were collected, and the fluorescent-expressing cell was sorted into each well of a 96-well plate containing a monolayer of MDCK cells. Following 48 h of incubation, a GFP-expressing or mCherry-expressing well was observed under a fluorescence microscope. The RNA of the fluorescent-expressing well was extracted to genotype the virions by Q-PCR. (B) Identification of the origin of genes from PR8-mCherry or VN-GFP viruses by MP-HRM. Curves colored red clustered with the PR8-mCherry, and curves colored green clustered with the VN-GFP. The expression of the reporter gene identified the origin of the NS gene. (C) MDCK cells were simultaneously infected with parental viruses at the indicated MOIs. At 12 h.p.i., cells were collected for flow cytometric analysis to determine co-infection frequency. Representative dot plots from infected MDCK cells assessed for co-infection.

### 2.3. Reassortment frequency enhanced with the increase of co-infection

Due to the segmented nature of the influenza genome, reassortment occurs when two influenza viruses co-infect one cell. To test the relationship of infection, co-infection, and reassortment, we performed a series of co-infections in MDCK cells with equal amounts (MOI) of PR8-mCherry and VN-GFP viruses at a range of multiplicities from MOI=0.1 (2^-3.32^) to 2^-12.32^. The cells were collected at 12 h.p.i. and flow cytometry was used to enumerate the infection rate. The supernatants were also collected to analyze the genetic constellation of the progenies further.

Co-infection was observed following co-inoculation of cells with virus stocks with a range from about 98.6% cells double infected at an MOI of 0.1 and about 0.27% at an MOI of 2^-12.32^ pfu/cell (Fig.2C and Fig.4A). Reassortment was detected in all the groups. The frequency of reassortant progeny enhanced with the increase of co-infected cells (the highest frequency [average 91.2%] at an MOI of 2^-5.32^ and the lowest frequency [average 32.2%] at an MOI of 2^-12.32^) (Fig.3 and Fig.4A). Even though percentage single positive cell, percentage double positive cell and percentage reassortment raised monotonically with the increase of inoculated dose, they showed differing patterns (Fig.4A). The percentage experimental reassortment was much higher than the percentage theoretical reassortment, which indicated that the semi-infectious particles might enhance reassortment. Besides, the % of fully infectious particles in the released progenies reduced with the increase of the inoculated dose from about 25% at an MOI of 2^-12.32^ to about 0.2% at an MOI of 2^-3.32^ (Fig.4B).

**Fig. 3.**
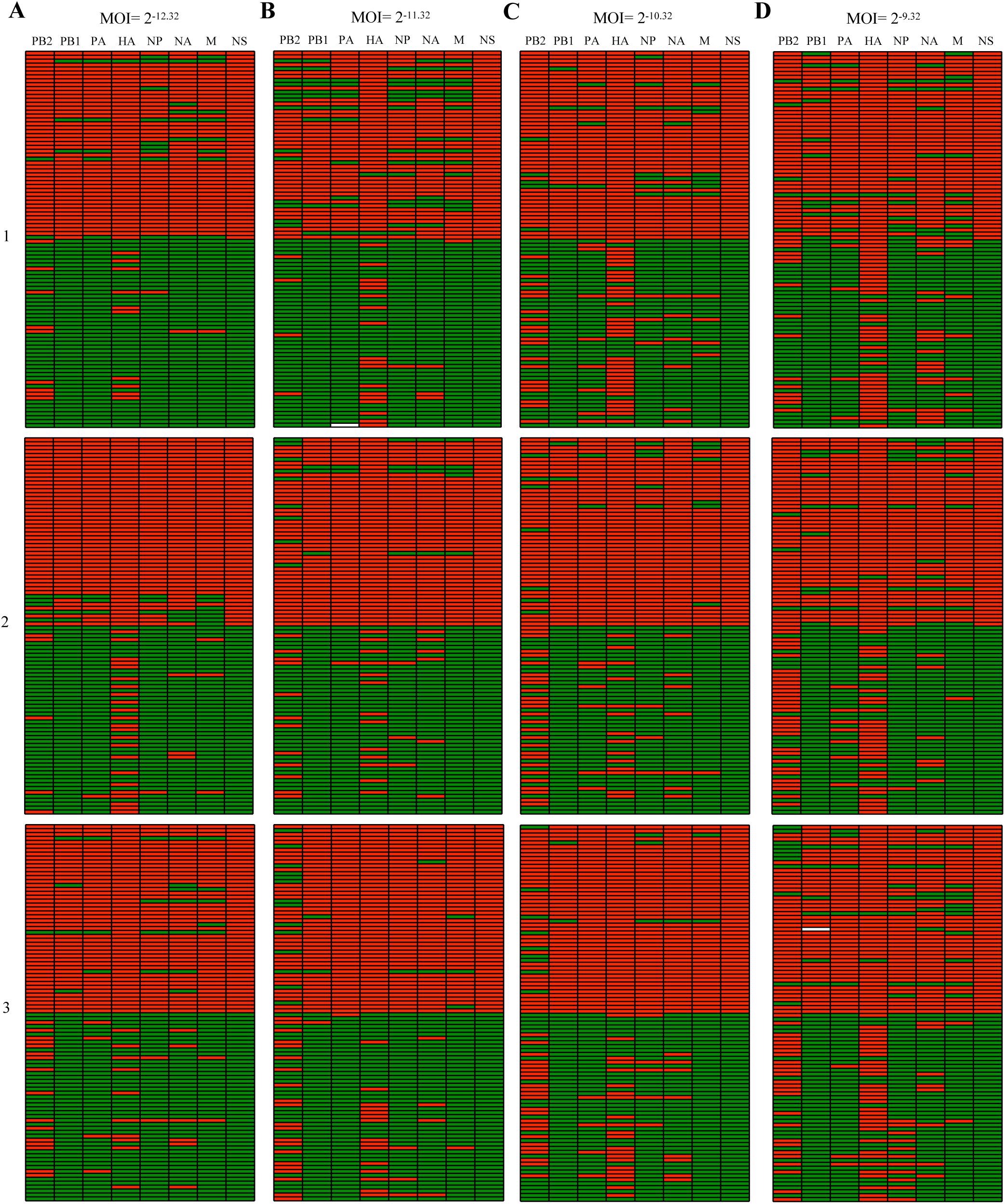
Enhanced reassortment frequency with the increase of co-infection. The results are shown of MP-HRM genotyping analysis of single-cell sorting progenies obtained from co-infected MDCK cells at the indicated MOIs. (A) 87/288 (30.2%) had reassortant genotypes at an MOI of 2^-12.32^. (B) 124/288 (43.1%) had reassortant genotypes at an MOI of 2^-11.32^. (C) 149/288 (51.7%) had reassortant genotypes at an MOI of 2^-10.32^. (D) 199/288 (69.1%) had reassortant genotypes at an MOI of 2^-9.32^. The genotype of individual progeny is depicted as rows in each panel with columns indicating the origin of each segment; red coloring indicates a segment derived from PR8-mCherry virus; green coloring indicates a segment derived from VN-GFP virus; white indicates a segment that was untyped.

**Fig. 4.**
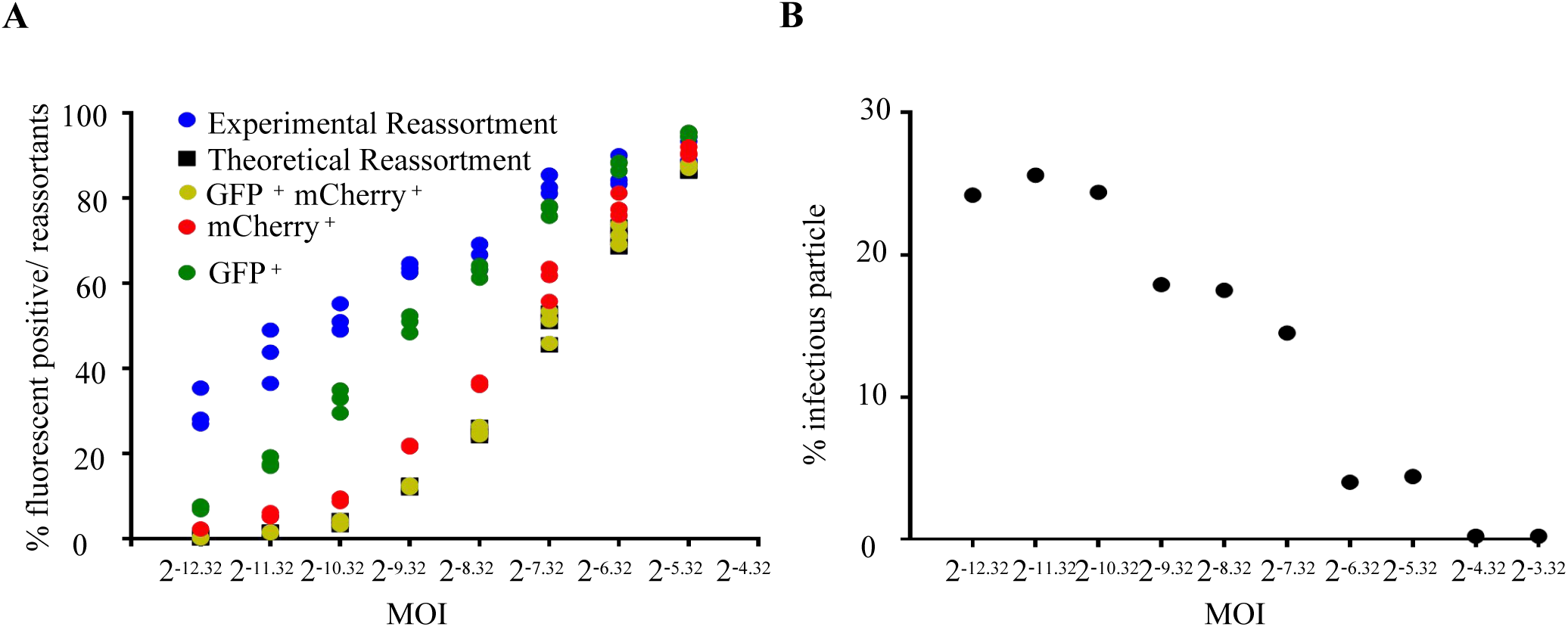
Enhanced reassortment rate but decreased infectious progeny rate with the increase of co-infection. (A) The frequency of reassortment enhanced with the increase of co-infection. PR8-mCherry and VN-GFP viruses were mixed in equal proportions and inoculated MDCK cells at a range of MOIs. Infection at each MOI was performed in triplicate. At 12 h.p.i., cell culture supernatants were stored, and fluorescent-positive cells were identified by flow cytometry. Progenies derived from each cell culture supernatant were isolated by single-cell sorting assay and genotyped by MP-HRM to measure % reassortment. Individual data points, each corresponding to one cell culture dish, are plotted. Red circles represent mCherry positive; green circles represent GFP positive; yellow circles represent double positive; blue circles represent experimental reassortment rate; black squares represent theoretical reassortment. (B) The percentage of fully infectious virions decreased with the increase of infection. The rate of fully infectious progeny was calculated according to the percentage of fluorescent-positive wells from single infected cells.

### 2.4. Strain-specific neutralizing mAbs shape antigenic shift during the co-infection *in vitro*

Reassortment is an essential evolutionary route for IAVs to generate pandemic strains [6–10]. This process is determined by various factors [11,13,16–27]. To understand the impact of cross-reactive immunity acquired via previous virus exposure on reassortment, we first tested the neutralization activities of three mAbs (6F12, directed against H1; GG3, directed against both H1 and H5; 1H4, directed against H5) when added to MDCK cells infected with PR8-mCherry and VN-GFP at an MOI of 2^-9.32^ of each virus. As previously reported, 6F12 only showed a high degree of neutralization activity to PR8-mCherry, 1H4 just showed efficacy against VN-GFP, and GG3 had vigorous neutralization activity against both parental viruses (Table 1 and Fig.6A) [31,41,42].

**Table 1.** Characterization of antibodies employed in this study.

| Name | Source | Specificity | Reactivity to H1 virus? | Reactivity to H5 virus? |
| --- | --- | --- | --- | --- |
| 6F12 | Mouse | HA Stalk | Yes | No |
| GG3 | Mouse | HA Stalk | Yes | Yes |
| 1H4 | Mouse | HA Head | No | Yes |

To evaluate the effect of mAbs on the reassortment, we then added the mAbs to co-infected (MOI=2^-9.32^ for each parental virus) MDCK cells at two h.p.i. For 6F12 and GG3, the expression of both parental NS genes was significantly reduced; 1H4 significantly prevented VN-GFP NS gene expression but not PR8-mCherry NS gene expression (Fig. 6B). These mAbs also significantly reduced the shedding of progeny virions in the supernatants of co-infected cells (Fig. 6C and D). To test the hypothesis that neutralizing mAbs promotes antigenic shift through reassortment, we analyzed the genotype of the progeny virions in the co-infected cells. The results showed that all these mAbs significantly reduced the release of progeny virions (Fig. 6C and D). Compared to the mock, the genotypes of the strain-specific mAbs were primarily changed. Notably, 6F12, which just neutralizes H1 viruses, promoted the proportion of H5 virus (both parental VN-GFP and reassortants bearing H5 HA) from 31.5% to 81.5%; 1H4, which only reacts with H5 viruses, increased H1 virus (both parental PR8-mCherry and reassortants bearing H1 HA) from 68.5% to 96.3%; GG3, which reacts with both parental viruses, did not obviously change the ratio of H1 viruses: H5 viruses (Fig.5 and Fig. 6E and F). Our results indicated that the addition of strain-specific neutralizing mAbs promote antigenic shift in co-infected cells.

**Fig. 5.**
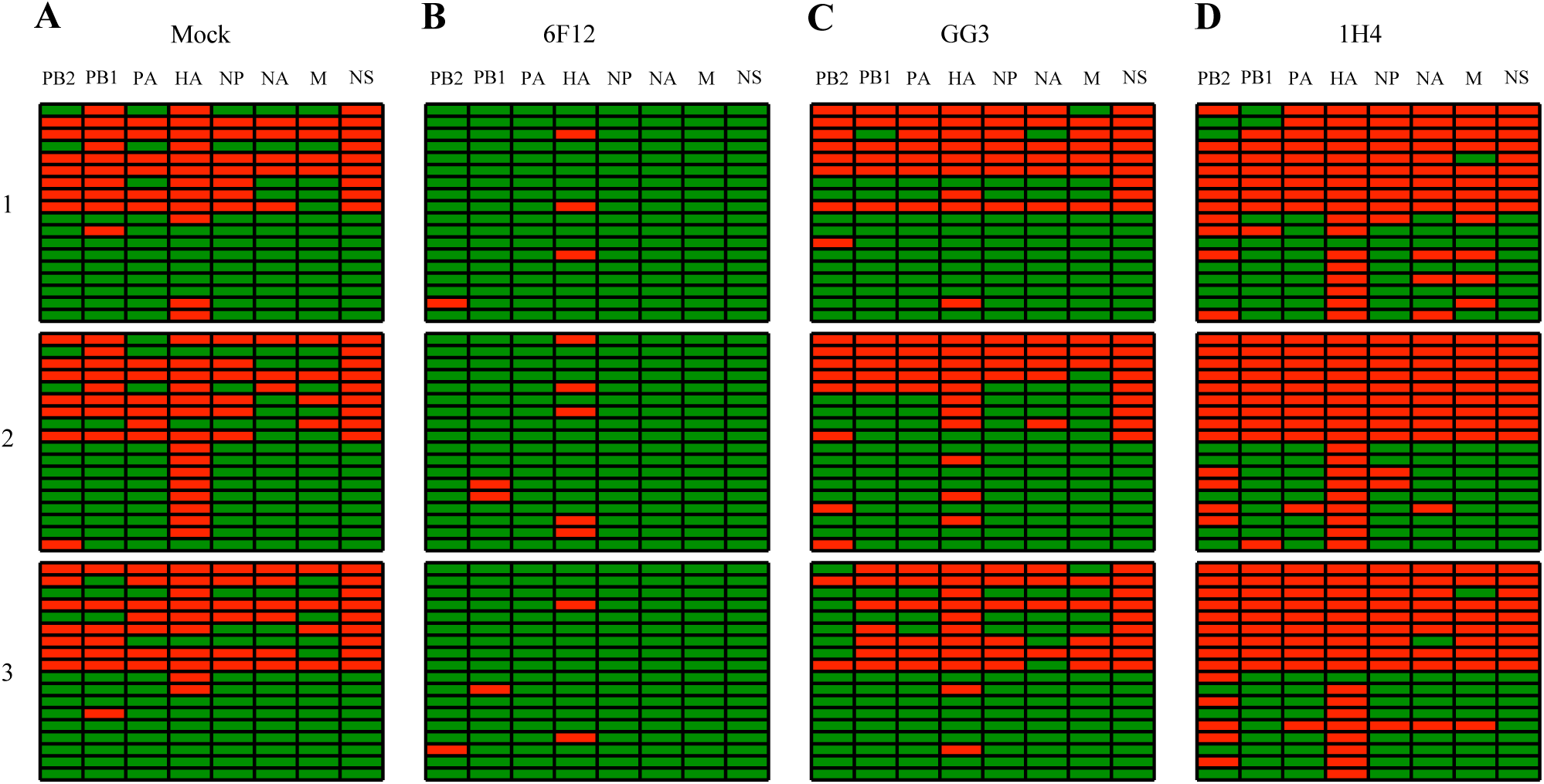
Neutralizing mAbs effecting reassortment in vitro. The results are shown of MP-HRM genotyping analysis of progenies obtained from co-infected MDCK cells with or without neutralizing mAbs. (A) MDCK cells were co-infected with parental viruses at an MOI of 2^-9.32^ of each virus, served as a negative control. (B) 30 ug/mL of mAbs 6F12 were added into the co-infected cells at 4 h.p.i.. (C) 10 ug/mL of mAbs GG3 were added into the co-infected cells at 4 h.p.i.. (D) 10 ug/mL of mAbs 1H4 were added into the co-infected cells at four h.p.i.. 54 individual fully infectious virions were genotyped in each group.

**Fig. 6.**
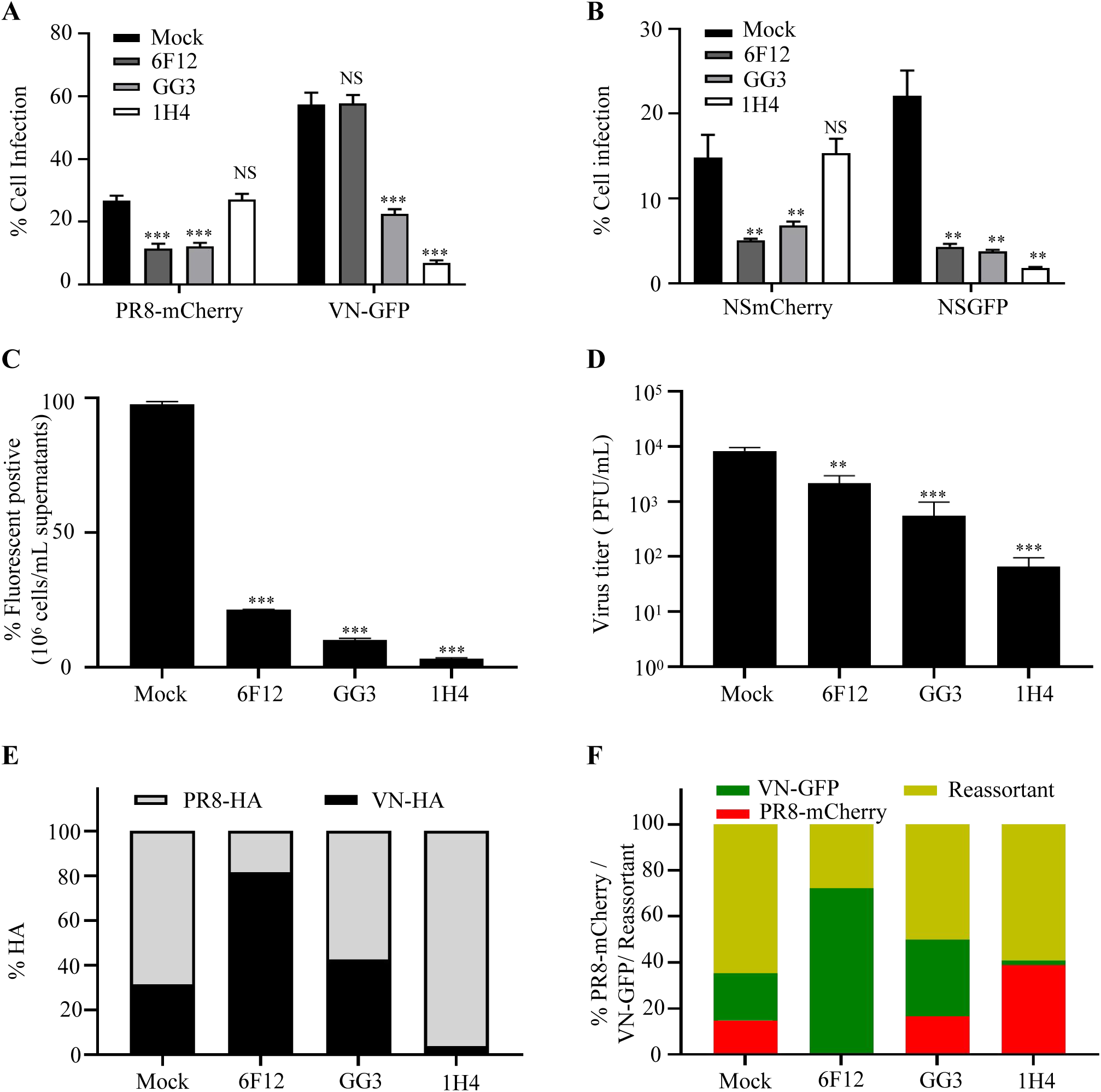
Strain-specific neutralizing mAbs shape antigenic shift during the co-infection in vitro. (A) Neutralizing ability of 6F12, GG3, and 1H4 mAbs against PR8-mCherry or VN-GFP virus. MDCK cells were infected with PR8-mCherry or VN-GFP virus at an MOI of 2^-9.32^. MAb (6F12, GG3, or 1H4) were added into the medium at four h.p.i.. The cells were harvested to calculate the infection rate. (B) Effect of neutralizing mAbs on the expression of NS genes in the co-infected MDCK cells. MDCK cells were co-infected with parental viruses at an MOI of 2^-9.32^ of each virus. MAb (6F12, GG3, or 1H4) were added into the medium at four h.p.i.. The cells were harvested to calculate the expression of NS genes, and the supernatants were collected for titration and genotyping. (C) Titration of the supernatants by flow cytometry. MDCK cells were infected with 2-fold serially diluted supernatants. At six h.p.i., the infection rate was quantified by flow cytometry. (D) Titration of the supernatants by plaque assay. (E) Effect of mAbs on HA gene in the progenies released from the co-infected cells. (F) Effect of mAbs on the reassortant rate in the progenies released from the co-infected cells. Data are shown as mean ± SD. One-way analysis of variance (ANOVA) test was used for statistical analysis. **, p < 0.01; ***, p < 0.001; NS, not significant.

### 2.5. Strain-specific neutralizing hemagglutinin-targeting monoclonal antibodies promote antigenic shift during the co-infection *in vivo*

To test how these neutralizing mAbs would impact reassortment and antigenic shift in an animal host, we performed prophylactic passive transfer challenge experiments in the mouse model (Fig. 8A). The study evaluated two key virological parameters: protection efficiency (quantified through survival rate analysis and viral titer measurement) and genetic reassortment (determined by virion genotyping). First, we performed a pre-mixed PR8-mCherry virus (2*10^5^ pfu) and VN-GFP virus (2*10^5^ pfu) challenge experiment with mice prophylactically receiving 3 mg/kg of the respective mAbs. All the mAbs were utterly protective, but the 1H4 group showed severe clinical symptoms and more weight loss (17.9%, 7 days post-infection (d.p.i.)). Animals pre-treated with 6F12 and GG3 showed no clinical signs with a weight loss of 3.8% and 2.7%, respectively, at seven d.p.i. (Fig. 8B and C). To investigate the reduction in lung virus titer, mice were pre-treated with 3 mg/kg of the respective antibody, challenged with pre-mixed viruses, and lungs were harvested on day 3 post-infection. All the neutralizing mAbs 6F12, GG3, and 1H4 significantly reduced lung titers compared to the mock group (Fig 8D and E).

To further assess the impact of antibodies on antigenic shift and reassortment in the co-infection system. Progeny virus isolates obtained from the lung samples were then genotyped. The results showed that all the mAbs affected the genotypes of the descendent virions. 6F12 suppressed the progeny viruses bearing H1 HA from 9.3% to 1.9%, and 1H4 decreased the proportion of H5 HA from 90.7% to 20.4% (Fig.7 and Fig. 8F and G), which indicated that strains-specific neutralizing mAbs promote an antigenic shift in the co-inoculated mice.

**Fig. 7.**
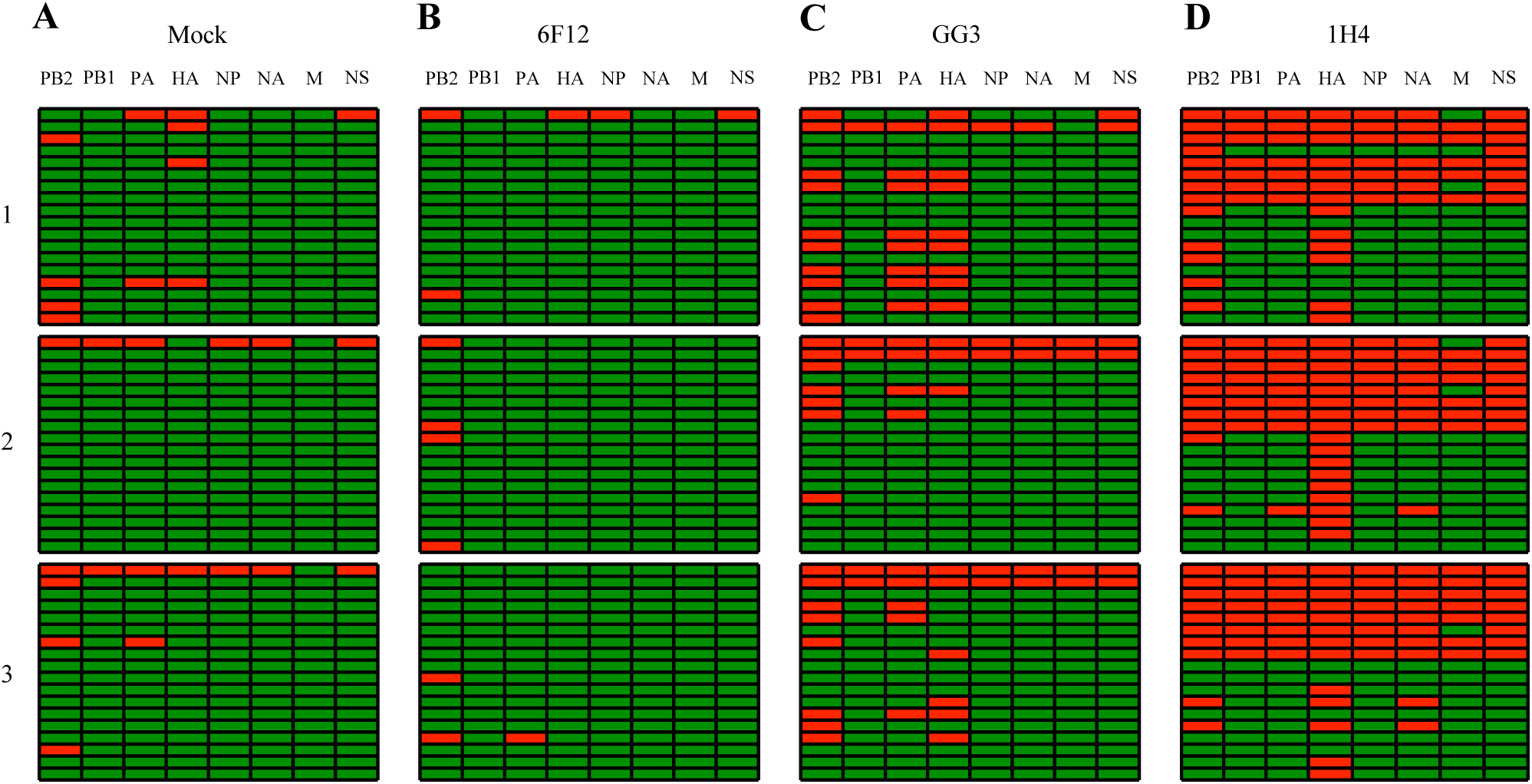
Neutralizing mAbs effecting reassortment in vivo. (A) Six- to 8-week-old BALB/c mice co-infected with parental viruses (2*10^5^ pfu of each virus), served as a negative control. Mice were intraperitoneally administered 3 mg/kg of body weight of mAb (B) 6F12, (C) GG3, or (D)1H4 two hours prior to challenge. The results are shown of MP-HRM genotyping analysis of fully infectious progenies obtained from lung samples at three d.p.i..

**Fig. 8.**
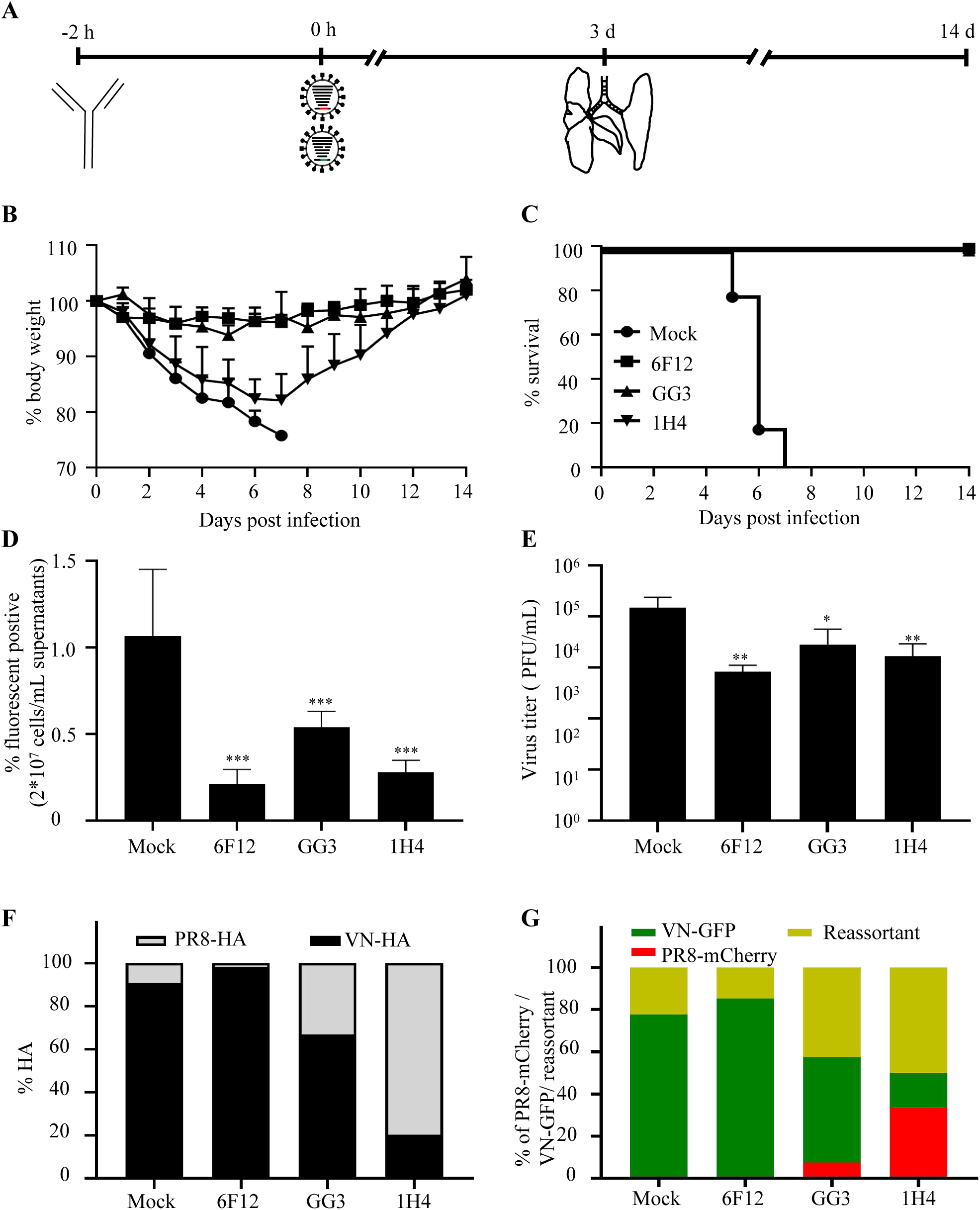
Strain-specific neutralizing mAbs shape antigenic shift during the co-infection in vivo. (A) Schematic representation of the passive immunization and challenge study. After co-infection, mice (n =5) were monitored daily for weight change (B) and survival (C). At three d.p.i., three mice from each group were randomly sacrificed, and lungs were harvested. Lungs were then homogenized to test viral lung titers by (D) flow cytometry or (E) plaque assay. (F) Effect of mAbs on HA gene in the progenies released from the co-infected mice. (G) Effect of mAbs on the reassortant rate in the progenies released from the co-infected mice. Data are shown as mean ± SD. One-way analysis of variance (ANOVA) test was used for statistical analysis. *, p<0.05; **, p < 0.01; *** p < 0.001.

## 3. Disscussion

Recent studies have elucidated the key factors governing the reassortment of influenza A viruses (IAVs)[12,13,15,16,19,43–46]. The emergence of pandemic strains, such as H2N2 and H3N2 in humans, and zoonotic IAVs, including H5N6, H5N8, and H7N9 in avian species, has been attributed to the selective pressure imposed by the host’s immune response[6,7,47–49]. However, the mechanisms underlying the relationship between immune pressure and reassortment remain poorly understood. To address this gap in knowledge, we developed a novel approach to investigate the effect of cross-immunity on reassortment events between two distinct IAV strains.

The segmented genome of influenza viruses is a major driver of their evolution [10]. Investigating the reassortment process between different influenza strains requires a versatile approach, involving at least four critical components: (i) robust labeling of parental viruses to track their origins; (ii) sensitive and reliable detection of co-infection events; (iii) unbiased isolation of progeny viruses to preclude experimental bias; and (iv) efficient and cost-effective techniques to assign the origin of each genome segment. Despite previous efforts to quantify influenza reassortment [13,50–53], existing methodologies have notable limitations. Here, we introduce three types of methods designed to overcome these shortcomings and enable precise, high-throughput analysis of reassortment in genetically diverse virus strains.

First, we developed two fluorescent-expressing influenza A viruses (IAVs), PR8-mCherry and VN-GFP, to facilitate real-time tracking of viral infections. Fluorescent IAVs are valuable tools for visualizing IAV-infected cells and have been described previously [36,38,54–57]. For example, Manicassamy *et al.* [36] used a fully replication-competent PR8-GFP IAV to monitor the dynamics of IAV infection *in vitro* and *in vivo*. Fukuyama *et al.* [38] made a panel of fluorescent IAVs to study the location and distribution of IAVs in whole-lung tissues of mice. Recently, fluorescent viruses have been made via tagging viral HA gene, which has significantly improved the investigation of reassortment [15,50]. However, this approach is more suitable for IAVs with identical HA genes and requires secondary labeling of infected cells. Moreover, fluorescent IAVs offer distinct advantages over other labeled viruses for the study of co-infection and reassortment, as they allow for real-time monitoring of viral infections. Our fluorescent IAVs, PR8-mCherry and VN-GFP, exhibited distinct phenotypes. Notably, PR8-mCherry was attenuated both *in vitro* and *in vivo*, whereas VN-GFP was not. Both fluorescent IAVs retained their pathogenicity in mice and stably expressed the reporter proteins after three successive passages in MDCK cells and mice. This finding is consistent with previous studies that demonstrated differential effects of reporters and virus backbones on virus attenuation, stability, and reporter activity [56,57]. Comparative analysis of co-infection rates revealed that our reporter virus system was more sensitive than previously reported methods [15]. Specifically, at an MOI of 0.1, the co-infection frequency of MDCK cells in our study was 99%, whereas the previous report showed a co-infection rate of less than 60%. The differential expression patterns of NS protein (intracellular) and HA protein (cell surface) or strain-specific differences may contribute to this disparity.

Second, we applied a single-cell sorting assay to isolate fluorescent-expressing cells to circumvent selection bias associated with variable progeny fitness. Traditional methods for isolating progeny virions from co-infected cells rely on plaque picking and purification [13,51–53]. However, this approach will lead to biased results, as progeny virions with varying fitness levels form plaques of different sizes and densities. Moreover, approximately 99% of influenza virions are noninfectious particles that can form plaques through complementation by co-infection [39,44], which further confounds the accuracy of reassortment measurements. To avoid these pitfalls, we infected cells with a mixed virion preparation at a low multiplicity of infection (MOI < 1), ensuring that fewer than 1% of cells were fluorescent-positive. At 6 h.p.i., we collected and sorted infected cells using a single-cell sorting assay. Notably, our results showed that higher infectious doses (MOI = 0.1) resulted in a lower percentage of fully infectious particles (0.2%) (Fig. 4B), highlighting the potential for contamination by noninfectious particles when isolating plaques. In contrast, our single-cell sorting approach enabled us to unbiasedly and accurately isolate fully infectious particles.

Third, the efficient genotyping of reassortant viruses requires labor-saving and cost-saving technologies. To date, several technologies have been developed to determine the origin of each segment. However, traditional methods such as polyacrylamide gel electrophoresis [51] and temperature-sensitive [52] approaches have constrained the selection of parental viruses. While PCR-based methods detected by gel electrophoresis can be used [15,24], they are labor-intensive. Additionally, although sequencing of PCR amplicons [13,53] and single-cell sequencing [58] are reliable approaches, their scalability is limited by high costs. As an alternative, High-Resolution Melt (HRM) analysis has been employed to differentiate between similar segments [15,50,59]. In this study, we designed a set of seven primer pairs to genotype the progeny virions, based on the conserved sequences of VN-GFP and PR8-mCherry. The NS gene was differentiated by the fluorescent protein in a single-cell sorting assay. This system enabled efficient quantification of the co-infection and reassortment of VN-GFP and PR8-mCherry in MDCK cells across a range of multiplicity of infection (MOI) values. Notably, the proportion of actual reassortant progeny viruses was significantly higher than the theoretical number calculated based on the co-infection rate, particularly at lower MOI values (Fig. 4A). This finding is likely due to the presence of semi-infectious particles that lack essential segments for labeled gene expression, thereby promoting reassortment through co-infection [44].

Reassortment is the outcome from a complex process. The fitness of reassortants, as well as the host’s selection pressures, influence the ability of more fit genotypes to become established in the host [9,10,15,60]. Specifically, the emergence of a reassortment virus is primarily driven by immune pressure, mainly in the form of antibody responses against the HA and NA proteins, acquired from prior exposure to historic virus strains through either vaccination or natural infection [13]. Two types of antibodies have been described to define the immune status against influenza viruses: strain-specific antibodies, which are specific to a particular virus strain, and cross-reactive antibodies, which can recognize multiple virus strains [29,34,41,42,61,62]. Here, we used three mAbs to investigate the role of previously acquired cross-immunity on reassortment. Tan *et al.* [29] previously reported that mAb 6F12 has neutralizing activity against a diverse panel of H1 viruses. However, for the parental viruses used in this study, 6F12 is a strain-specific mAb, capable of neutralizing PR8-mCherry but not VN-GFP. Similarly, 1H4 is a strain-specific mAb that targets VN-GFP [41]. In contrast, GG3 is a cross-reactive antibody that can neutralize both parental viruses [63]. Our *in vitro* co-infection results showed that both strain-specific neutralizing mAbs and cross-reactive neutralizing mAbs significantly reduced the release of progeny virions (Fig. 6C and D). Notably, the addition of mAbs led to the detection of completely different genotypes of virions compared to mock-treated cells, indicating that prior immunity plays a crucial role in shaping viral reassortment in MDCK cells. Specifically, the HA gene of the progeny viruses was significantly affected by the strain-specific neutralizing mAbs compared to the cross-reactive antibody.

Previous studies have shown that vaccination protects pigs against challenge from co-inoculated viruses, although different reassortant viruses have been detected in co-infected pigs with and without preexisting immunity [13]. As previously reported, the antibodies used in this study conferred protection in an antibody transfer challenge study in mice [29,42]. However, mice developed divergent clinical symptoms and weight loss, suggesting that the strain-specific variations in genotype correlated with diverse pathogenicities. Consistent with our *in vitro* findings, strain-specific neutralizing mAbs significantly influenced the reassortment-driven production of virions bearing non-neutralized HAs. Our results indicate that preexisting immunity drives the evolution of IAVs.

In summary, we developed and characterized fluorescent-expressing IAVs as a tool for measuring reassortment. Using this approach, we established a sensitive and unbiased assay to quantify the co-infection and reassortment of antigenically diverse IAVs. We then used neutralizing monoclonal antibodies (mAbs) to examine the influence of pre-existing immunity on reassortment. Our findings showed that strain-specific neutralizing mAbs selectively promote the generation of reassortant viruses bearing non-neutralized HA segments, indicating that these antibodies drive antigenic shift during reassortment. Ultimately, our findings highlight a critical concern for vaccine design, as the mismatch between existing immunity and circulating IAV strains can inadvertently accelerate the emergence of novel pandemic viruses.

## 4. Materials and Methods

### 4.1 Cell culture

MDCK cells and Human embryonic kidney (293T) cells were maintained in Dulbecco’s minimum essential medium (MEM) supplemented with 10% fetal bovine serum. The cell reagents were purchased from Gibco Life Technologies.

### 4.2 Plaque assay

MDCK cells were seeded in a 6-well plate format with a density of 1×10^6^ cells per well. The following day, the MDCK cells were washed twice with PBS and incubated with 200 μL of opti-MEM containing 10-fold serial diluted virus for one h at room temperature with frequent shaking. The cells were washed twice with PBS and then overlayed with MEM containing a 2% oxoid agar and 1 μ g/mL of L-(tosyl amido-2-phenyl) ethyl chloromethyl ketone (TPCK)-treated trypsin (Sigma). At 48 h post-infection, the cells were fixed with 4% paraformaldehyde (PFA) for one h, and the overlays were removed. The plaques were visualized by staining with Crystal Violet.

### 4.3. Generation of NS-fluorescence virus

#### 4.3.1 Construction of NS-fluorescence segment

The mCherry virus constructs were cloned using the NS gene segment of PR8. The GFP virus constructs were cloned using the NS gene segment of VN, and the mult-basic cleavage site of HA was removed to reduce virulence.

As previously described, the NS segment fused with the reporter gene was constructed by overlapping fusion PCR [36,63]. In brief, we modified the NS segment to express NS1-reporter and NEP as a single poly-protein with a 19 aa porcine teschovirus-1 (PTV-1) 2A autoproteolytic cleavage site between them (Fig. 1A). Additionally, silent mutations were introduced to the endogenous splice acceptor site in the NS1 ORF to prevent splicing [64]. The constructed NS segment was subsequently inserted into the pDZ IAV rescue plasmid [38,65].

#### 4.3.2 Rescue of NS-fluorescence virus

The viruses (PR8, PR8-mCherry, VN, VN-GFP) used in this study were generated using standard reverse genetics techniques [37,64]. Briefly, 0.5 μg of each of the eight pDZ plasmids representing the eight segments of the IAV genome were transfected into 293T cells using Lipofectamine 2000 (Invitrogen). After 24 h, the 293T cells were re-suspended in the cell medium, and 100 μL of the mixture was injected into 8 to 10-day-old embryonated eggs. The virus was then harvested from the allantoic fluid at 48 h.p.i.. The successful rescue of the virus was confirmed by conducting a hemagglutination assay with chicken red blood cells. After three passages in MDCK cells, the fluorescent-expressing plaques were picked to make a stock in MDCK cells. The sequence of the eight segments in the virus was verified by sequencing. The viral titer was determined by performing plaque assay in MDCK cells.

#### 4.3.3 Single-cycle and multi-cycle growth curve

MDCK cells were seeded at a dilution of 10^6^ cells/well in 6-well plates 12 hours before infection. The cells were washed twice with PBS, infected with viruses at MOI of 1 and 0.001, incubated for one hour on ice with frequent shaking, washed three times with PBS, and then cultured with Opti-MEM/1 μ g/mL TPCK-treated trypsin. Virus titers were determined by plaque assay in MDCK cells at the indicated time points.

#### 4.3.4 Stability of the parental viruses *in vitro*

To test the stability of the parental reporter viruses *in vitro*, MDCK cells were infected with PR8-mCherry or VN-GFP at an MOI of 0.001. Supernantant were collected at 24 h.p.i.. MDCK cells were inoculated with 200 μ L of opti-MEM containing three times 10-fold diluted virus supernatant from the previous MDCK passage. The viruses were blindly passaged thrice in MDCK.

### 4.4 Co-infection at a range of MOIs with two parental viruses in MDCK cells

Each well of a 6-well plate was seeded with 1 × 10^6^ MDCK cells 12 h before infection. PR8-mCherry and VN-GFP viruses were diluted to 10^6^ pfu/mL in PBS and mixed in a 1:1 ratio. Each virus mixture was then diluted with PBS to the appropriate titer for inoculation at MOI 2^-3.32^, 2^-4.32^, 2^-5.32^, 2^-6.32^, 2^-7.32^, 2^-8.32^, 2^-9.32^, 2^-10.32^, 2^-11.32^, or 2^-12.32^ pfu/cell of each virus. Before infection, the growth medium was removed, and the monolayer cells were washed with PBS twice. Then, each well was inoculated with a 200 µL volume and incubated on ice for one hour to ensure equal binding of all viruses. After 1 hour, the unattached virus was removed by washing three times with PBS. Subsequently, the virus medium (opti-MEM supplemented with 1 μg/mL trypsin) was added, and the cells were transferred to 37℃. After 12 hours post-infection, the supernatant was collected and stored at -80 ℃ for subsequent genotyping of the released virus using the single-cell sorting assay. The infected cells were harvested and prepared for flow cytometry.

### 4.5 Flow cytometry

#### 4.5.1 Measurement of % infection, % co-infection

To determine the % infection and % co-infection, MDCK cells were harvested 12 h after co-infection with PR8-mCherry/VN-GFP viruses by using TPCK-treated trypsin. The cells were washed thrice with PBS and re-suspended in 500 µL PBS containing 2% BSA. Flow cytometry was performed using a BD FACS Aria III SROP (Beckman Coulter), and analyzed with Flow Jo software [19].

#### 4.5.2 Virus titration

Flow cytometry and the plaque assay are two methods used to titrate the co-infected samples (supernatants of cells and supernatants of the homogenized lung of mice).

To choose an appropriate infectious dose of the co-infected samples for the single-cell sorting assay (see below), virus titration was performed by flow cytometry based on the detection of mCherry/GFP expressed in infected cells. Briefly, the supernatants were serially diluted in 2-fold dilutions in PBS. The MDCK cells in 6-well plates were washed twice with PBS and incubated with 200 μL of virus for 1 hour on ice with frequent shaking. After washing with PBS three times, 2 mL of opti-MEM medium was added. The cells were transferred to 37℃ and were harvested at six h.p.i.. Viral infection was then quantified by flow cytometry. Besides, traditional plaque assay was also utilized to titrate the co-infected samples.

### 4.6 Single-cell sorting assay

MDCK cells were infected with supernatants collected from co-infected samples at a low MOI (less than 1% of cells will be fluorescent-expressing), incubated on ice for one h, washed three times with PBS, incubated at 37℃ for six h, harvested with trypsin, and washed three times with PBS. The single fluorescent-expressing cell was then sorted into each well of a 96-well plate containing a monolayer of MDCK cells. Following 48 h of incubation, a GFP-expressing or mCherry-expressing well was observed under a fluorescence microscope and selected for genotyping the virions.

### 4.7 Determination of virus genotypes by high-resolution melt analysis

To genotype the fully infectious virions isolated by single-cell sorting assay, After incubating for 48 h, RNA of the fluorescent-expressing well was extracted with the viral RNA kit. 4 μL of RNA were reverse transcribed using the Prime Script ™ RT Master Mix reverse transcriptase (Takara) following the manufacturer’s instructions. The resulting cDNA was then used as a template in qPCR reactions. Specifically, four μ L of 1:4 diluted cDNA was combined with the appropriate primers (0.4 μM final concentration, primer sequences shown in Table S1) and the TB Green®Premix Ex Taq ™ II kit (Takara). qPCR and melt analyses were carried out using a Roche480 II Real-Time PCR Detection System.

### 4.8 Impact of mAbs on co-infection and reassortment in MDCK cells

Murine mAbs 6F12 (directed against the H1 stalk domain) [39], GG3 (directed against the H1 and H5 stalk domain) [42], and 1H4 (directed against the H5 head domain) [41]were produced from hybridomas previously generated using a classical hybridoma fusion protocol [41].

To determine the effect of pre-existing mAbs on co-infection and reassortment, MDCK cells in a 6-well plate format were washed twice with PBS and subsequently co-infected with 2^-9.32^ pfu/cell of each parental virus. Single virus infection (PR8-mCherry or VN-GFP) was conducted as a control. One hour post incubation on ice, cells were washed three times with PBS and incubated in opti-MEM at 37°C with 5% CO_2_ for four hours. After that, the medium was removed, and the cells were cultured for 8 hours at 37°C in opti-MEM containing trypsin and the appropriate concentration of mAb (6F12, 30 ug/mL; GG3 10 ug/mL; or 1H4, 10 ug/mL). After eight hours of incubation with mAbs, the supernatants were harvested to determine the effect of mAbs on reassortment frequency, and cells were collected to calculate the effect of mAbs on co-infection.

### 4.9 Mouse experiments

Animal procedures followed protocols approved by the Animal Care and Use Committee (ACUC) at Inner Mongolia University (SYXK 2020-0006). All animal experiments were performed in the negative-pressure isolators of the authorized animal biosafety level 2 (ABSL-2) facility. Six- to eight-week-old female BALB/c mice were obtained from SPF (Beijing) Biotechnology Co., Ltd (Beijing, China). Mice were anesthetized by Pentobarbital before infection. Virus was given at the indicated amounts diluted in 50 µL volume of phosphate buffered saline (PBS) via the intranasal route, equally divided over both nares. Animals body weight conducted by researchers unaware of treatment conditions to eliminate observer bias in mortality determination were monitored daily for 2 weeks. For lung virus titration, mice were euthanized at the indicated time points in the text and lungs were removed aseptically and homogenized in PBS before being flash frozen in liquid nitrogen and stored in -80 degrees C until titration by plaque assay on MDCK cells.

Sample sizes were determined by assay requirements: three mice per group for viral stability/genotyping (to balance consensus accuracy with 3R principles), and five mice per group for MLD50 and antibody protection (ensuring 80% power to detect large effects, based on pilot data SD=20%). This is consistent with previous virology studies.

#### 4.9.1 Determination of MLD_50_

The MLD_50_ was determined for selected groups of viruses representing four different levels of pathogenicity, namely PR8, PR8-mCherry, VN, and VN-GFP. 100 BALB/c mice were randomly distributed into 4 treatment groups (25 mice/group), with each group further divided into 5 cages of 5 mice. Ten-fold serial dilutions were prepared using a sterile solution of PBS and antibiotics, ranging from 10^7^ to 10^1^ pfu. Each mouse was i.n. infected with 50 µL of the diluted virus. A group of 5 BALB/c mice were inoculated with each virus after being anesthetized. Following the virus challenge, the mice were monitored randomly for 14 days post-infection for disease symptoms such as scruffy fur, hunched appearance, reduced bright-alert response, weight loss, and mortality. According to our animal protocol, mortality was recorded based on actual death or a 25% weight loss cut-off. The MLD_50_ values were calculated as EID_50_ using the Reed-Muench method [66].

#### 4.9.2 Stability of the parental viruses *in vivo*

To evaluate the stability of the parental reporter viruses *in vivo*, a total of 18 BALB/c mice were randomly allocated into 2 equal groups with three subgroups of 3 mice each. We administered intranasally (i.n.) to 3 mice with a dose of 10^4^ pfu of PR8-mCherry or VN-GFP. After three d.p.i., the lungs of each mouse were collected and homogenized (10% w/v) in PBS. Then, the supernatant of lung homogenate was centrifuged. The other three naive mice were administered i.n. with the lung homogenate from previously infected mice. The process of i.n. inoculation of three BALB/c mice was repeated 3 times. The percentage of fluorescent-expressing plaques measured the viruses’ stability in cells and mice.

#### 4.9.3 Passive transfer experiments in mice

To evaluate the effect of pre-existing mAbs on reassortment, mouse passive transfer experiments were performed. Four groups of BALB/c mice (n=8 per group) were established through randomization of 32 total animals. Mice were intraperitoneally treated with 3 mg/kg of body weight of mAb 6F12, GG3, 1H4, or PBS. After a 2-hour treatment, the mice were i.n. co-infected with 2*10^5^ pfu of each virus (PR8-mCherry and VN-GFP).

Groups of five mice were randomly monitored daily for clinical signs of illness and weight loss throughout the 2-week experiment. Upon reaching 75% of initial body weight, animals were humanely euthanized with Pentobarbital as per the ACUC protocol.

Groups of three mice were euthanized on 3 d.p.i.. The lung was collected to determine virus titers and analyze the genotype of the progeny virions on MDCK cells.

### Statistical information

Statistical analyses were performed using the Prism 9.0 program (GraphPad, San Diego, CA, USA). Data were compared using One-way and Two-way analysis of variance (ANOVA) tests for statistical analysis. * p<0.05, **p < 0.01, *** p < 0.001. NS is not significant.

## Acknowledgements

This project was supported by a grant from the Inner Mongolia Autonomous Region Major Science and Technology Project (2021ZD0013), the National Key Laboratory of Reproductive Regulation and Breeding of Grassland Livestock (Jointly Built by the Province and Ministry) - Identification of Specific Target Points for Important Pathogens in Cattle and Sheep and Development of Novel Diagnostic Technologies (2025KYPT0066), the ‘Grassland Talents’ Program of Inner Mongolia Autonomous Region (12000-12102617), the High-Level Talents Research Support Program of Inner Mongolia Autonomous Region (10000-21311201/004), the ‘Horse Program’ High-Level Talents Program of Inner Mongolia University (10000-21311201/141), , the Science and Technology Leading Talent Team Program of Inner Mongolia Autonomous Region (2022LJRC0009 to W. Hu from Inner Mongolia University and Fudan University), the Science and Technology Major Special Project of Inner Mongolia Autonomous Region (2020ZD0008 to Prof. Wei Hu from Inner Mongolia University and Fudan University). G.W. began this work as a postdoctoral researcher in Adolfo García-Sastre’s lab at the Ichan School of Medicine at Mount Sinai. We sincerely thank Dr. García-Sastre for supplying the plasmids for influenza virus rescue and the murine mAbs used in this study.

## Author contributions

G. J.W., R.H. and H.W. conceived and designed the experiments; G.J.W., M.N.L, X.R.G., X.Y.L., Y.T., S.Q.W., J.Y.H. and W.Y.Y. performed the experiments; G.J.W., R.H. M.N.L and X.R.G. analyzed the data; G.J.W. and R.H. interpreted the data; G.J.W. and M.N.L wrote the original draft; G.J.W., C.L.and R.H. reviewed and edited the paper.

## Materials & Correspondence

Correspondence and requests for materials should be addressed to Guojun Wang.

## Supplemental material

**Table S1 Primers used to generate amplicons for high-resolution melt analysis**

