## Supplemental Table 1 for "Subtype-specific neutralizing antibodies promote antigenic shift during influenza virus co-infection": Table S1.pdf

Table S1 Primers used to generate amplicons for high-resolution melt analysis

| Segment | Forward primer | Reverse primer |
|---|---|---|
| PB2 | 477F<br>CCATGCAGATCTCAGTGCTAAA | 1584R<br>TTTCTCTGTTCCCTGGGTTTC |
| PB1 | 2016F<br>TCTTCCCCAGCAGTTCATA | 2319R<br>AGTAGAAACAAGGCATTTTTTCA |
| PA | 1864F<br>AACAAATCAGAAACATGGCC | 2077R<br>TCAATTGCTTCATATAGCCC |
| HA | 1584F<br>GGAGTRAAATTGGAATCAAT | 1755R<br>AGTAGAAACAAGGGTGTTTTT |
| NP | 1213F<br>CCAGAAGYGGAGGAAACACC | 1544R<br>AGTAGAAACAAGGGTATTTTTC |
| NA | 1167F<br>GATTGGTCAGGRTATAGCGG | 1389R<br>AGTAGAAACAAGGAGTTTTTTGAAC |
| M | 159F<br>GGCTAAAGACAAGACCAATCCT | 574R<br>CTCCATAGCCTTAGCTGTAGTG |
| NS | 1626F<br>CAGAAGTTTGAAGAAATAAG | 1756R<br>AGTAGAAACAAGGGTGTTTTTTATC |
